# Efficient 3D cone trajectory design for improved combined angiographic and perfusion imaging using arterial spin labeling

**DOI:** 10.1101/2024.01.17.576043

**Authors:** Qijia Shen, Wenchuan Wu, Mark Chiew, Yang Ji, Joseph Woods, Thomas W Okell

**Author notes:** Please address correspondence to: Prof Thomas Okell WIN/FMRIB Centre, John Radcliffe Hospital, Headley Way, Headington, Oxford, OX3 9DU, UK.

## Abstract

**Purpose:** To improve the spatial resolution and repeatability of a non-contrast MRI technique for simultaneous time resolved 3D angiography and perfusion imaging by developing an efficient 3D cone trajectory design.

**Methods:** A novel parameterized 3D cone trajectory design incorporating the 3D Golden Angle was integrated into 4D combined angiography and perfusion using radial imaging and arterial spin labeling (CAPRIA) to achieve higher spatial resolution and sampling efficiency for both dynamic angiography and perfusion imaging with flexible spatiotemporal resolution. Numerical simulations and physical phantom scanning were used to optimize the cone design. Eight healthy volunteers were scanned to compare the original radial trajectory in 4D CAPRIA with our newly designed cone trajectory. A locally low rank reconstruction method was used to leverage the complementary k-space sampling across time.

**Results:** The improved sampling in the periphery of k-space obtained with the optimized 3D cone trajectory resulted in improved spatial resolution compared with the radial trajectory in phantom scans. Improved vessel sharpness and perfusion visualization were also achieved in vivo. Less dephasing was observed in the angiograms due to the short echo time of our cone trajectory and the improved k-space sampling efficiency also resulted in higher repeatability compared to the original radial approach.

**Conclusion:** The proposed 3D cone trajectory combined with 3D Golden Angle ordering resulted in improved spatial resolution and image quality for both angiography and perfusion imaging and could potentially benefit other applications that require an efficient sampling scheme with flexible spatial and temporal resolution.

## Introduction

Visualizing the dynamic process of blood flow along the vascular tree to the tissue facilitates the diagnosis and treatment monitoring of various cerebrovascular diseases. Stroke, for example, is a significant cause of mortality and disability and is caused by a reduction in blood supply, either from blocked/narrowed arteries (ischemia) or bleeding into the brain tissue (hemorrhage)^1^. Similarly, moyamoya disease is caused by the progressive narrowing of intracranial arteries, leading to downstream tissue ischemia^2^. Arterial spin labeling (ASL) is a non-invasive, non-contrast based imaging technique^3,4^ that can be used to generate angiographic and perfusion images. However, the acquisition time for separate angiographic and perfusion scans may be too long for busy clinical settings. A recently designed acquisition method, 4D Combined Angiography and Perfusion using Radial Imaging and ASL (CAPRIA),^5,6^ provides a scheme to simultaneously map the dynamics of blood flow through the arteries and at the level of the brain tissue, enabling both sets of information to be obtained in a time-efficient manner. High spatial/temporal resolution angiograms and low spatial/temporal resolution maps of tissue perfusion can be reconstructed from the same raw CAPRIA data. Yet, the resolution and SNR of the images is constrained by the limited sampling efficiency of the radial trajectory used by the sequence, with poorer sampling density in the periphery of k-space and the requirement for many excitation pulses, which attenuate the ASL signal. Improved sampling density at the edges of k-space and increased k-space coverage per excitation are desirable to improve both spatial resolution and SNR. This will enable more accurate diagnosis with improved visualization of small distal vessels and subtle perfusion alterations and motivates our search for a more efficient k-space trajectory for CAPRIA that maintains its flexibility in spatiotemporal resolution.

In addition to radial trajectory, numerous other designs of 3D non-Cartesian trajectories have been proposed over the years. Stack-of-stars^7^ and stack-of-spirals^10^ were proposed to improve sampling efficiency but were anisotropic between slice and within slice. Stack-of-cones^9^ was designed to uniformly cover k-space in a shorter total scan time than an equivalent Cartesian trajectory, while satisfying the Nyquist sampling theorem. Density adapted radial^10^, variable density cones^11^ and FLORET^12^ are some variations of the radial and cone trajectory with improvements in sampling efficiency and uniformity. Other trajectories, such as radial-cones^13^ and Seiffert spirals^14^, focus on incoherent and relatively uniform sampling, but the Nyquist sampling theorem is no longer strictly enforced. Optimization-based trajectories like SPARKLING^15^, PILOT^16^ and SNOPY^17^ are gaining more attention recently, but the optimal trajectory usually takes a time-consuming process to recalculate for specific scanning parameters.

To improve on 4D CAPRIA, we require a trajectory that: i) can be simultaneously optimized for both angiography and perfusion imaging; ii) samples a large region of k-space with each readout to minimize the number of excitation pulses required, and therefore the attenuation of the ASL signal; iii) has a short echo time to maximize SNR and minimize flow-based dephasing; iv) accommodates dynamic imaging with variable temporal resolution; and v) has different but comparable sampling patterns for each temporal frame that result in incoherent aliasing across time to enable the use of temporal regularization in the reconstruction. Unfortunately, although they have many desirable features, none of the trajectories outlined above satisfy all of these requirements. The anisotropic sampling of stack-of-cones creates difficulty in undersampling different temporal frames with comparable sampling patterns. Radial-cones specifies the direction of each shot with isotropically distributed radial directions, which is not suitable for varying the temporal resolution post-hoc. Additionally, the trajectory formulation does not provide a means of trading off the sampling density in peripheral and central k-space, which are more important for angiography and perfusion imaging, respectively.

In this work, we aimed to improve the spatial resolution and SNR for angiographic and perfusion images acquired with 4D CAPRIA. Taking inspiration from the radial-cones approach^13^, we present an efficient and flexible two-stage parametric cone design that can be combined with golden ratio rotations around the k-space center to satisfy the trajectory requirements outlined above. We then used numerical simulations and physical phantom scanning to optimize the cone shape parameters. Finally, we performed in vivo scans with healthy volunteers to demonstrate the superior image quality obtained with our cone trajectory compared to the original radial approach.

## Methods

### Pulse Sequence Design

As illustrated in Figure 1, the original 4D CAPRIA method^6^ consists of a pre-saturation module (to remove spin history effects and reduce static tissue signal), a pseudocontinuous ASL (PCASL) ^18^ labeling preparation (to label/control blood magnetization as it flows through a defined labeling plane in the neck) and a continuous golden ratio radial readout to image the blood as it traverses the arterial tree (allowing the generation of angiographic images) and exchanges into the tissue (yielding perfusion images). In this implementation, an additional inversion pulse is added to improve the suppression of static tissue signal, reducing physiological noise, as per^19^. The blood signal is isolated from static tissue signal by subtracting control and label data. High spatiotemporal resolution angiograms can be reconstructed using a small temporal window, giving a high undersampling factor (∼50X undersampled) that can be tolerated due to the sparse and high SNR nature of the angiographic signal. Lower spatiotemporal resolution perfusion images can be reconstructed from the same raw data using a broader temporal window and only using the samples in a central region of k-space (∼4X undersampled), improving the conditioning of the reconstruction for this weaker and less sparse signal. Variable flip angles are used in the train of readouts to reduce the signal attenuation from the initial excitation pulses and boost the perfusion signal at later timepoints.

**Figure 1.**
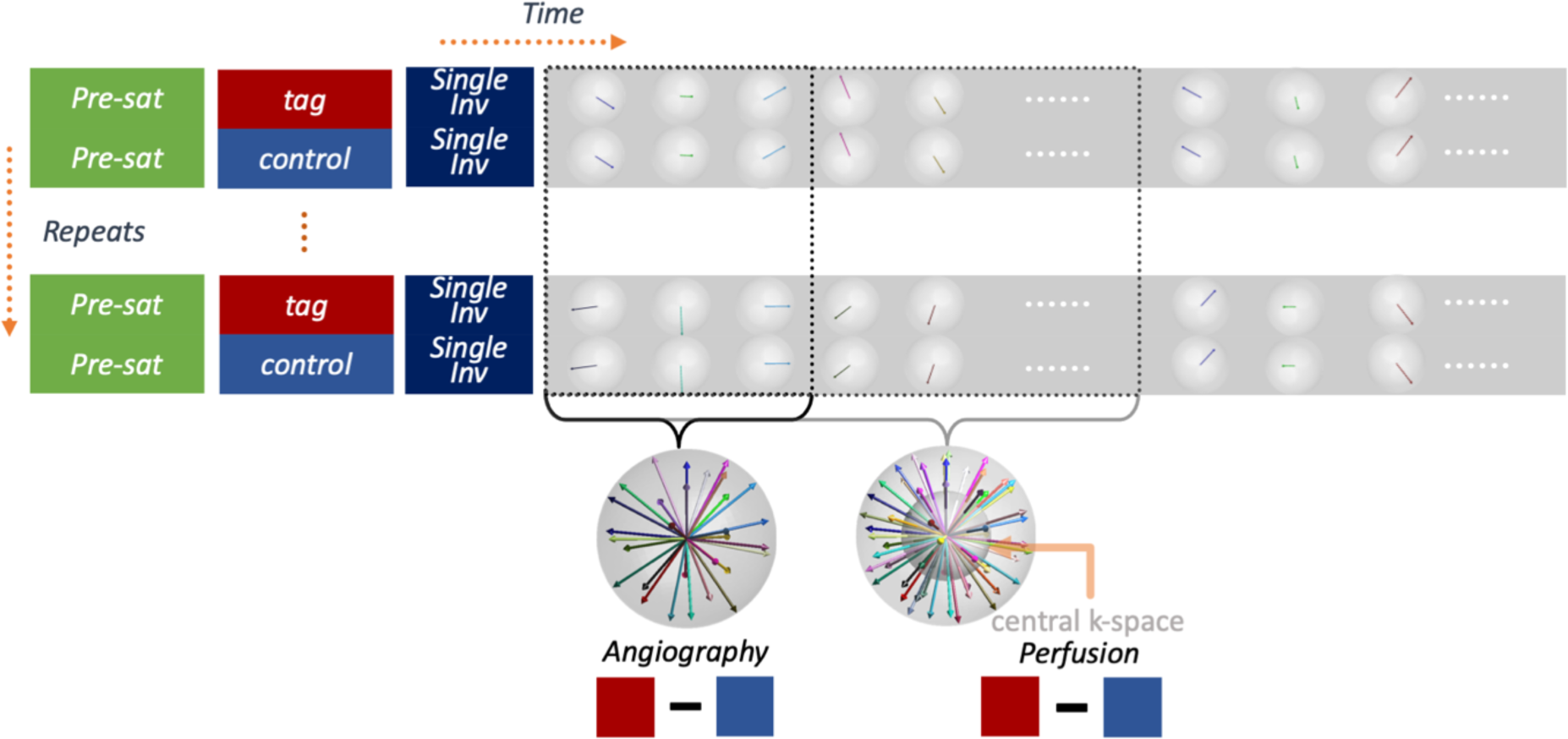
Schematic pulse sequence diagram. The preparation period (pre-saturation, pseudo-continuous ASL labeling and a slab-selective inversion pulse) is followed by a continuous golden ratio readout. The radial case is shown here, representing the original 4D CAPRIA approach, but the proposed cones case is similar, with each radial spoke replaced by a cone at the same orientation. Samples across all repeats can be flexibly combined across time with variable temporal resolution, whilst maintaining approximately uniform sampling. Examples for grouping one frame of angiography (high spatiotemporal resolution) and perfusion imaging (low spatiotemporal resolution, using only the central region of k-space) are illustrated using red and blue boxes, respectively.

**Table 1.** Acquisition parameters for the three different in vivo protocols.

| protocol name<br>Acquisition parameters |  | Optimal radial protocol | Matched radial protocol | Cone (Ours) |
| --- | --- | --- | --- | --- |
| **Label** | Labeling duration | 1.8 s |  |  |
| **Readout** | Flip angle | 2~9° | 3~12° | 3~12° |
|  | readout time | 6.6 ms | 10 ms | 10 ms |
|  | TR | 9.8 ms | 14.7 ms | 14.7 ms |
|  | TE | 4.56 ms | 6.26 ms | 1.35 ms |
|  | # of readouts after each ASL preparation | 216 | 144 | 144 |
|  | Acquisition time | 6 min 12 s | 6 min 12 s | 6 min 12 s |

CAPRIA relies on grouping readouts from multiple repeats for the reconstruction of both sets of images. In this work, we use the golden ratio ordering approach proposed by Song et al.^20,21^, incrementing the golden ratio rotations first across repeats and then across time, to maintain exact golden ratio ordering when grouping readouts across repeats for any temporal resolution. The original 3D radial approach is flexible, approximately isotropic, and motion robust, making it suitable for this application. However, the limited coverage of each radial spoke limits its k-space sampling efficiency.

### Base cone trajectory design

Inspired by 3D cones trajectories ^9,11,13^, in this work each CAPRIA radial spoke is replaced by a cone to give a much broader coverage of k-space after each excitation pulse. The excellent sampling efficiency of the cone is integrated with the advantages of a 3D golden ratio radial trajectory, to improve the spatial resolution and SNR while keeping the temporal flexibility for dynamic imaging.

A two-stage parametric design is proposed that jointly considers the sampling density within central and peripheral regions, because: 1) angiographic reconstructions utilize the full k-space data, whereas perfusion reconstructions only use the central region of k-space; and 2) spending more time in central k-space could improve robustness against motion and other sources of signal variation, whereas more samples in peripheral k-space could help improve spatial resolution. This is done by combining a conventional cone shape with a fixed cone angle^9^ for the central region of k-space (stage 1) with a cone whose cone angle increases with the radial distance for the peripheral region of k-space, as in radial-cones^13^(stage 2). The overall shapes of two parts for a base cone oriented along the k_z_ axis are described by Equation 1.

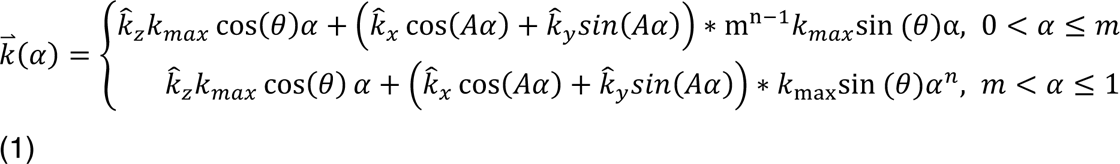

Where 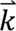 represents the trajectory coordinates in 3D k-space; 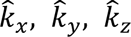 are three orthogonal unit vectors; θ, *m*, *n* are three free parameters that enable adjustment for different sampling requirements: θ is the cone angle spanned by the endpoint of the trajectory and the central axis, *m* is the separation point for the two parts, and *n* controls the curvature for the cone trajectory (as shown in Figure 2c); α describes the relative position along the trajectory; and *A* is a scaling factor to accommodate an arbitrary trajectory duration and gradient hardware limitations. Example trajectories with various parameter choices are shown in Supplementary Figure 1b,c.

**Figure 2.**
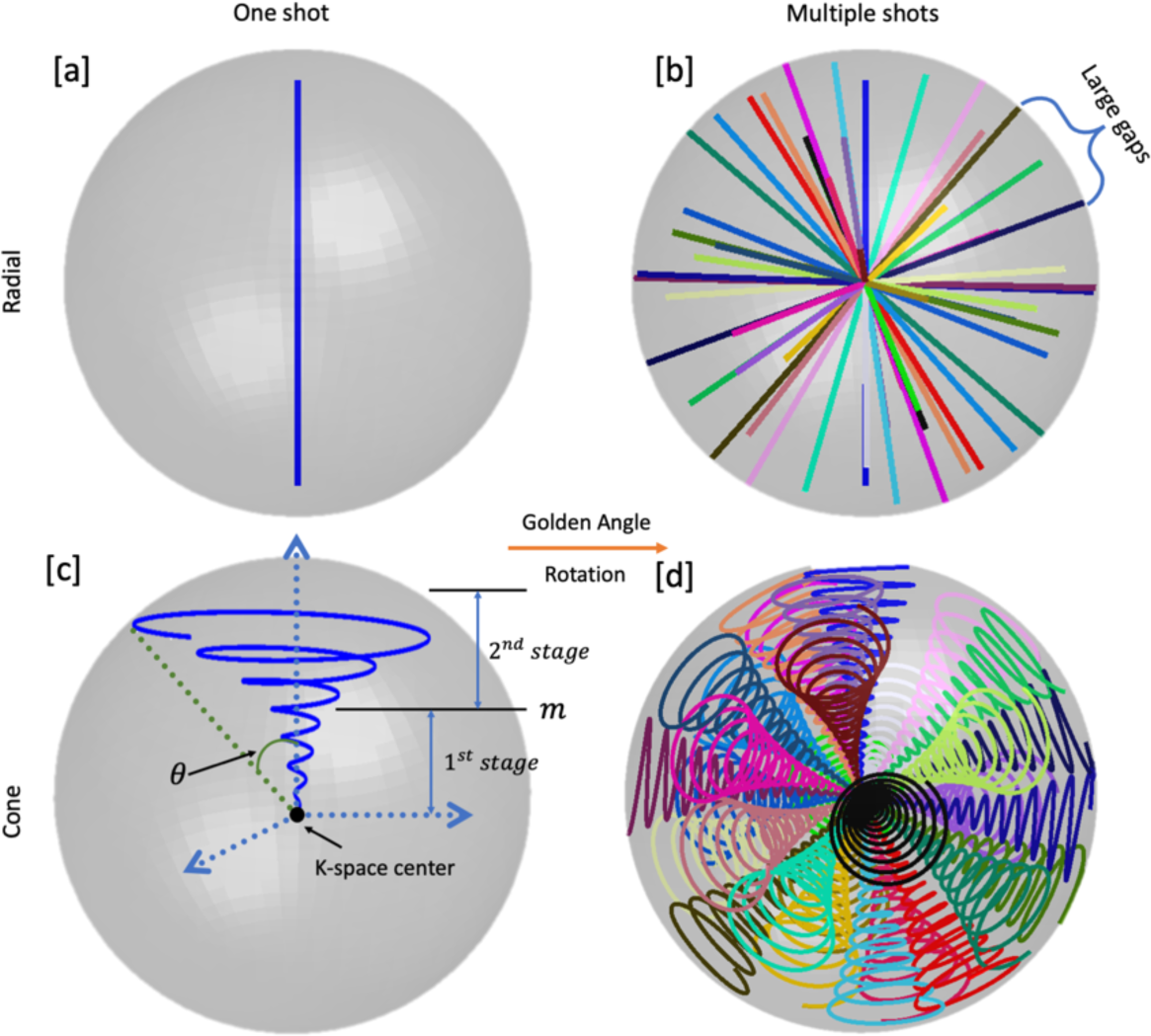
Design of the cone trajectory and the rotation scheme. Exemplary single radial spoke used in the original 4D CAPRIA is plotted in [a]. [b] demonstrates 30 radial spokes, each rotated using the 3D golden ratio relative to the previous spoke. Our cone design is detailed in [c], with shape control parameters cone angle θ, two-stage separation m in the plot. [d] is obtained by replacing the radial spokes in [a,b] by the cone trajectory in [c]. The comparison between [b] and [d] emphasizes the advantages of cone trajectory in terms of sampling efficiency and k-space coverage, particularly in the periphery.

Once θ, *m*, *n* and the readout duration have been chosen, the gradient waveforms and k-space samples for a given system performance (maximum gradient amplitude and slew rate) can be solved using a constrained optimization, as in Lustig et al.^22^. To reduce peripheral nerve stimulation and gradient imperfection effects, in our trajectory, the maximum gradient amplitude and slew rate were limited. (i.e., *G_max_* = 11.64 *mT* / *m*, *S_max_* = 61.54 *T/m/s*).

### Cone rotations

Given a single base cone trajectory (referred to as one shot below), the base shape can be rotated around the k-space center to cover the sphere of k-space to be sampled. As illustrated in Figure 2c,d, the central axis is first rotated using the 3D golden ratio ^23^. The cone is further rotated around its central axis by an additional random angle. This random rotation provides sufficient incoherency and uniformity without loss of flexibility for altering the total number of shots as shown in Supplementary Figure 1d. The golden ratio approach specifies a series of sequential acquisition directions so that any set of readouts with contiguous golden ratio ordering covers k-space approximately uniformly.

### Optimization of cone parameters

Unlike numerically optimized approaches, the proposed trajectory has only a few free parameters that need to be optimized. In preliminary experiments it was found that metrics derived from numerical simulations alone could only partially capture the imaging performance in real experiments due to complex interactions between signal attenuation, aliasing patterns, spatial resolution and robustness to real-world off-resonance effects and signal variations. Therefore, the free trajectory parameters were jointly optimized based on the results of numerical simulations and phantom scanning, utilizing the efficiency of numerical simulations and fidelity of physical phantom scans.

First, a wide range of parameters *m*, *n*, θ and readout time were simulated and compared. Using these simulations, the candidate parameter values were narrowed down to a smaller set of combinations, which were then further investigated with phantom scans to explore the behavior of the different trajectories under the effects of eddy-currents, off-resonance, and signal variability.

### Numerical simulations

For all simulation experiments, we simulated a single acquisition frame for both angiographic and perfusion images. A previously acquired high quality angiography image^24^ and a simulated perfusion phantom^25^ were used as the ground truth for all trajectories. The reconstructed spatial resolution was 1.13 *mm* and 3.39 *mm* isotropic for angiography and perfusion images, respectively. The number of repeats in sequence was fixed to 48 in all experiments, so the total number of shots is dependent on the temporal window of each binned frame. Perfusion images were reconstructed using all shots but only the central region of k-space 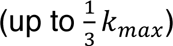. Angiographic images used the full k-space trajectory, but only half the temporal window (half of the shots). The numerical ground truth of both modalities is shown in Supplementary Figure 2.

The range of cone parameters used in simulations were as follows:

– the power term *n*: 1, 2 *or* 3.
– The separation point *m* for the two-stage design: 0 (i.e., single stage) or 0.5 (i.e., two-stage, transition of cone shape halfway through the trajectory)
– the cone angle θ: 15°, 30°, 45°, 60° *or* 75°

As resulting trajectory would not vary greatly with slight change of parameters, discrete values were tested in experiments.

Three metrics were calculated for comparison, each characterizing different aspects of the trajectory in this highly undersampled case:

1. Full width half maximum (FWHM): characterizes the spatial resolution by measuring the width of the central peak of the point spread function (PSF)
2. Effective SNR: characterizes how the thermal noise in the acquisition is transformed into noise in the output image. The pseudo-replica method^26^ was used, such that reconstructions were performed by adding many different instances of complex random Gaussian noise to the k-space data. The standard deviation of the reconstructed images across noise instances was taken as the “noise” and the mean value within the foreground (where the ground truth image has non-zero intensity) as the “signal” to calculate the SNR. Since spatial resolution also affects SNR, this effect was compensated by dividing the calculated SNR with the product of the FWHM in three orthogonal planes (approximating the true voxel volume).
3. Signal to aliasing power ratio (SAPR). This metric evaluates non-Cartesian artifacts in the background relative to the signal in the foreground. In numerical simulation, the average power of reconstructed image *I* within foreground mask *M_f_* (where ground truth image has non-zero intensity) was divided by that within background mask *M_b_*(voxels not included in the foreground). *N_f_*, *N_b_* represents thenumber of voxels within foreground and background masks respectively.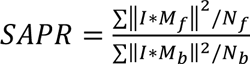

### Experiments: Physical phantom acquisition

Following on from the numerical simulations, in the physical phantom scans *m* and *n* were fixed to 0.5 and 3 respectively, and the effect of the cone angle θ and readout time were further investigated within the following ranges:

– The cone angle ϕ: 30°, 45° and 60°

– The readout time: 6.6 *ms*, 8 *ms* and 10 *ms*

A quality assurance phantom (GE DQA Phantom; GE Healthcare) was used to visualize spatial resolution changes and artifacts caused by the interaction of signal variations with different non-Cartesian sampling patterns. Additionally, only a single frame was acquired, reconstructed, and analyzed using the density compensated adjoint NUFFT because the temporal stability of the trajectory ensured by the 3D golden ratio means that subsequent frames would have similar properties.

### Experiments: In-vivo scanning protocol

Based on the numerical simulation and phantom experiments, a combination of the 4 parameters (*m*, *n*, θ, *readout time*) was selected that resulted in high SNR and spatial resolution and few artifacts (see Results section). These parameters were used for the cone trajectory in vivo with 4D CAPRIA.

To assess the proposed cone trajectory, it was compared with radial trajectories using two different protocols: the first was a protocol optimized in a previous study ^6^(referred to as optimal radial protocol) while the other matched the TR and flip angle schedule of the cone protocol (referred to as matched radial protocol), to ensure any differences seen were not attributed solely to the changes in these other parameters. In total, 8 subjects (age range 25-35) were scanned on a 3T Prisma scanner (Siemens Healthineers, Erlangen, Germany) using a 32-channel receive-only head coil.

A quadratically varying flip angle scheme was separately optimized for the different protocols using the same method as the original 4D CAPRIA sequence^6^. For the optimal radial protocol, the flip angles ranged between 2° − 9°. For both the matched radial protocol and the cone protocol, the flip angles ranged between 3° − 12°. The temporal resolution used to reconstruct perfusion and angiographic images was 352.8 *ms* and 176.4 *ms*, respectively. Note that the total time for all readouts after each ASL labeling module (2116.8 *ms*) as well as the total scan time (6 *min* 12 *s*) were kept consistent between different protocols for a fair comparison.

Coil compression^27^ was performed, reducing the original 32 channel data to 8 channels for reconstruction. Coil sensitivity maps were estimated from a pseudo-structural image (reconstructed by regridding averaged label and control data from the final 352.8 *ms* of the readout where the static tissue signal is more consistent) using the adaptive combine method.^30^ K-space data were multiplied by a Hann window prior to reconstruction to reduce undersampling artifacts.

A T_1_-weighted MPRAGE image with 1.5 *mm* isotropic resolution was acquired for each subject for analysis (described below).

### Image reconstruction

In numerical simulation and phantom scanning, the adjoint NUFFT^29^ was used to reconstruct every image for a faithful reflection of the SNR and spatial resolution using different sampling patterns.

For reconstructions of in vivo data, a locally low rank (LLR) regularization^30^ was used to exploit the spatiotemporal redundancy in the data and the complementary sampling at different time points, in a similar manner to previous work^6^:

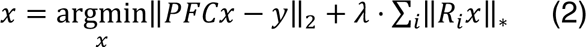

Where *x* is the image to reconstruct, *y* is the acquired k-space data, *P* is the sampling mask, *F* is the Fourier encoding matrix, *C* are the coil sensitivity maps, λ is a regularization factor and *R_i_* is the operator extracting each patch (indexed by i) from each frame. The patch size and regularization parameter were chosen empirically: λ = 10^’(^and patch size 5 × 5 × 5.

### Mean repeatability calculation

For in vivo experiments, a repeatability metric was calculated to report the scan-rescan stability. Specifically, each subject was scanned using the three protocols described above, with each protocol being repeated twice back-to-back to minimize motion or other signal variation between repeated scans.

To minimize the impact of relative motion between repeated scans, a pseudo-structural image was reconstructed for each scan by taking the mean of label and control signals, which were then registered to the MPRAGE image using affine registration (FSL’s FLIRT^31^). The same linear transformation was then applied to angiography and perfusion images. We then followed the steps described in Okell et al.^6^ to calculate scan-rescan repeatability for angiography and perfusion images separately. The results were Fisher transformed prior to performing the multi-way ANOVA on the group level with all 8 subjects to determine whether differences due to trajectory, timepoint or subject were significant.

## Results

### Numerical simulations

Quantitative results from the numerical simulations used to optimize the cone trajectory are shown in Figure 3. For brevity, only the most important metrics (FWHM and effective SNR) are displayed for angiographic (full k-space, temporal resolution of 176.4 *ms*) and perfusion (central k-space with radius 1/3 k_max_, temporal resolution of 352.8 *ms*) reconstructions, with the rest of the metrics and comparisons shown in Supplementary Figures 3-4. In Figure 3a-b, our two-stage design (*m* = 0.5) is compared with the single-stage design (*m* = 0), similar to radial cones^13^. Despite some variations across different cone angles (θ), the two-stage design (*m* = 0.5) consistently outperformed the single-stage design (*m* = 0) for both angiographic and perfusion reconstructions, in terms of spatial resolution and SNR. Similar results were obtained using readout times of 8 *ms* and 6.6 *ms* (see Supplementary Figures 3-4).

**Figure 3.**
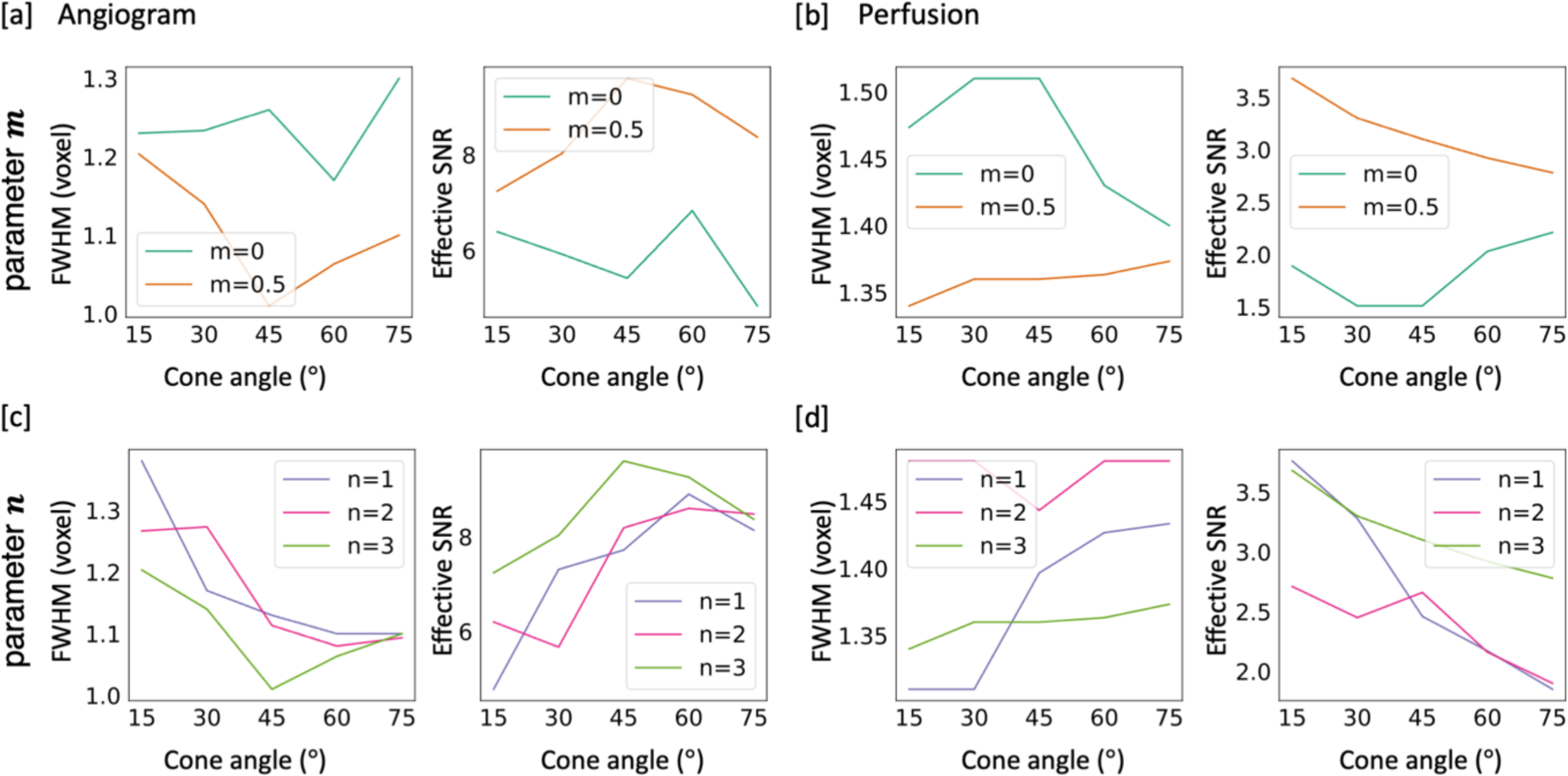
Representative numerical simulations for determining m and n in the cone design. In [a, b], readout time was fixed to 10 ms and n was fixed to 3. In [c,d], readout time was fixed to 10 ms and m was fixed to 0.5. [a] Simulation of angiography images to compare different selections of m on a range of cone angles. [b] Simulation of perfusion images using the same set of trajectories as [a]. Note that two-stage option m=0.5 surpassed the conventional single stage design (m=0) in resolution and SNR for both modalities regardless of cone angles. [c,d] Simulation for the effects of parameter n on angiography and perfusion images respectively, demonstrating that n=3 generally results in better spatial resolution and effective SNR. For angiography [a,c], the cone trajectory achieves the best resolution and effective SNR when the cone angle falls within the range 30°∼60°. For perfusion imaging [d], these metrics do not have a strong cone angle dependence (note the y-axis scaling).

When varying the parameter (*n*) that controls the curvature of the cone trajectory (Figure 3c-d), both the FWHM and the effective SNR fluctuate across different cone angles, but the trajectory generally performs best when *n* = 3. Both FWHM and effective SNR metrics are optimal at a cone angle of around 30∼60° in the case of angiography, with a weaker dependence for perfusion imaging. Note that the small perturbation in these metrics across different parameter choices depends on the complex interactions and overlap between neighboring cones, so there are no simple monotonic relationships here. A combination of *m* = 0.5 and *n* = 3 was chosen based on these numerical simulation results to narrow down the parameter space to be investigated using a physical phantom.

### Phantom scanning

Representative results of phantom scanning are shown in Figure 4, with the remaining comparisons in Supplementary Figure 5-6. Figure 4a shows the comparison between radial and cone trajectories with cone angles of 30° and 60°. Consistent with numerical simulations, the radial images are much blurrier than the cone images, due to the sparser sampling of peripheral k-space. In addition, radial trajectory produced coherent streaking artifacts(indicated with the red arrow). 60° cone trajectory, however, generated images with better discernable thin structures (pointed by the blue arrows) and superior SNR (zoomed in regions) compared with 30° cone and radial trajectories.

**Figure 4.**
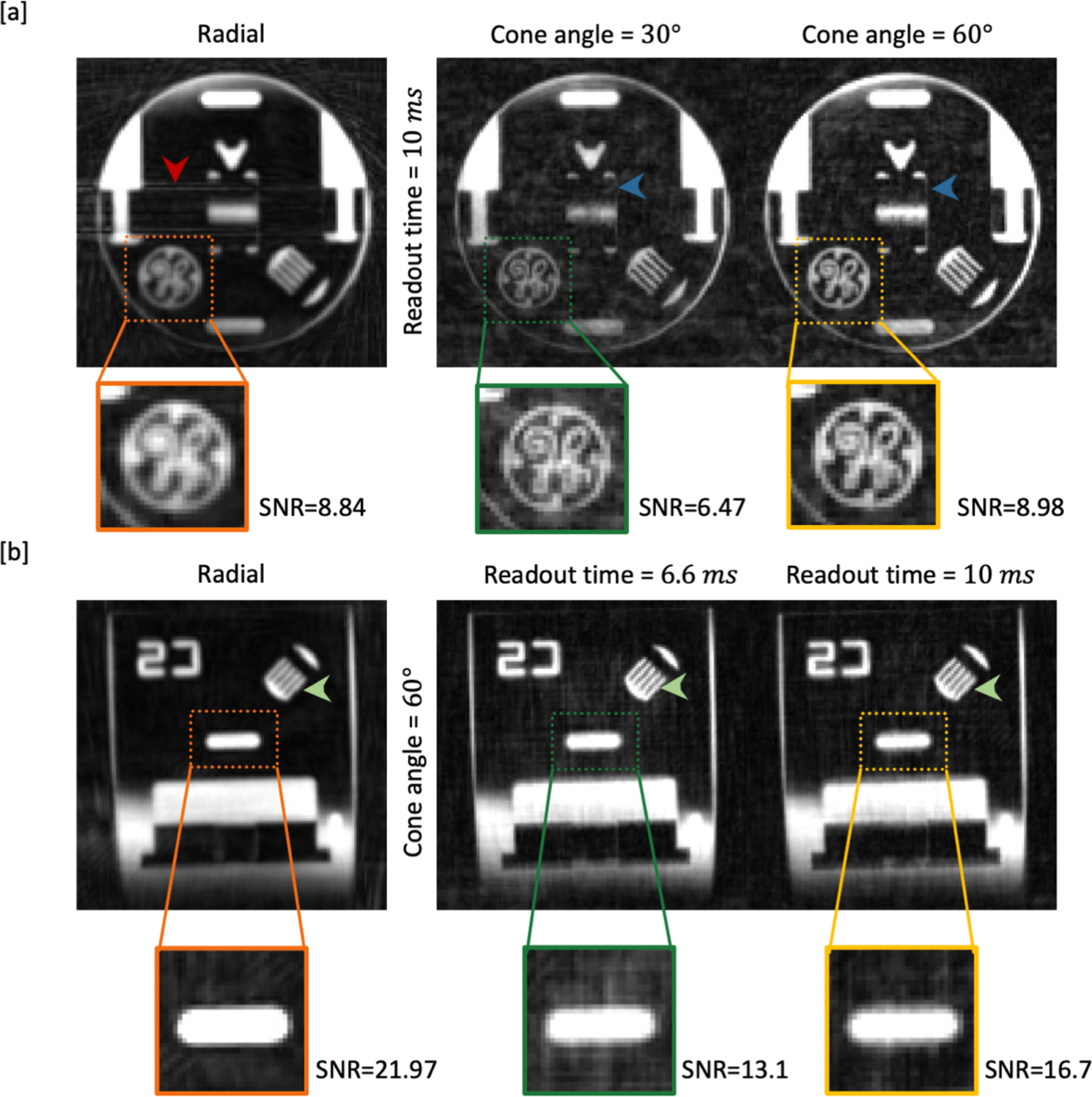
Comparison of a physical phantom scanned with different protocols and reconstructed with a density compensated adjoint NUFFT. [a] Here the readout time was fixed to 10 ms while the cone angle was varied between 30°∼60°. Undersampling in the radial trajectory resulted in coherent streaking artefacts (red arrow), whereas in the cone trajectories aliasing was more noise like, but less prominent in the 60° cone design, allowing fine details to be visualized (blue arrows). [b] Here the cone angle was fixed to 60° while the readout time varied between 6.6 ms ∼ 10 ms. The improved visualization of thin structures demonstrated the better spatial resolution of both cone protocols compared to radial (green arrows). Among the cone protocols, the lowest level of background artefacts was achieved when the readout time was 10 ms and the cone angle was 60°. More examples are given in supplementary Figures 5-6. The SNR was also reported on the zoomed in patches. A foreground mask was first extracted from the radial result by thresholding at 10% of maximum intensity. All voxels not included in the mask were considered as background. The same mask was used to calculate SNR on all trajectories in [a] and [b] respectively.

In Figure 4b, the cone angle is fixed to 60° while the readout time is varied. The comb-like structure pointed to by the green arrow further confirms the improved resolution using the proposed cone trajectory. Here, the SNR of cone trajectory with longer readout time surpassed that of shorter readout times, while the best SNR of radial trajectory was mainly attributed to its intrinsic smoothing with the wider point spread function.

Combining the results of the numerical simulation and the physical phantom scanning, using a cone trajectory with a larger cone angle (60°) helps improve spatial resolution and reduce background signal due to aliasing. A longer readout time is also favorable for reducing artifacts and has the additional advantage that fewer excitations are required after each ASL preparation, reducing the attenuation of the ASL signal and allowing higher flip angles to be used to further increase the SNR. Two factors prevent us from choosing a very long readout time. Firstly, the undersampling factor could be too high to compensate from longer readout time when the TR is too long, as we need to keep the total readout time after each labeling module fixed. Reducing the number of shots could also increase the anisotropy of the binned k-space coverage. Secondly, off-resonance issues become more prominent with longer readouts. Therefore, in this work, a combination of 10 *ms* readout time with 60° cone angle was chosen for the in-vivo acquisition.

### In-vivo perfusion results

Figure 5 presents a comparison of in vivo perfusion images acquired using radial and cone trajectories in one subject. Figure 5a compares the proposed cone protocol (readout time 10 *ms*, flip angles 3-12°) with the previously optimized radial protocol^6^ (readout time 6.6 *ms*, flip angles 2-9°) as well as a radial protocol with a readout time and flip angle schedule matched to that of the cone protocol (readout time 10 *ms*, flip angles 3-12°). Qualitatively, the cone trajectory demonstrated considerable improvement in visualizing fine tissue structures, whereas the radial results suffered from blurring that obscured some of these details. In the case of radial trajectory, the matched protocol with a longer TR and fewer spokes tended to yield more pronounced blurriness due to the limited peripheral k-space coverage and more T2* decay.

**Figure 5.**
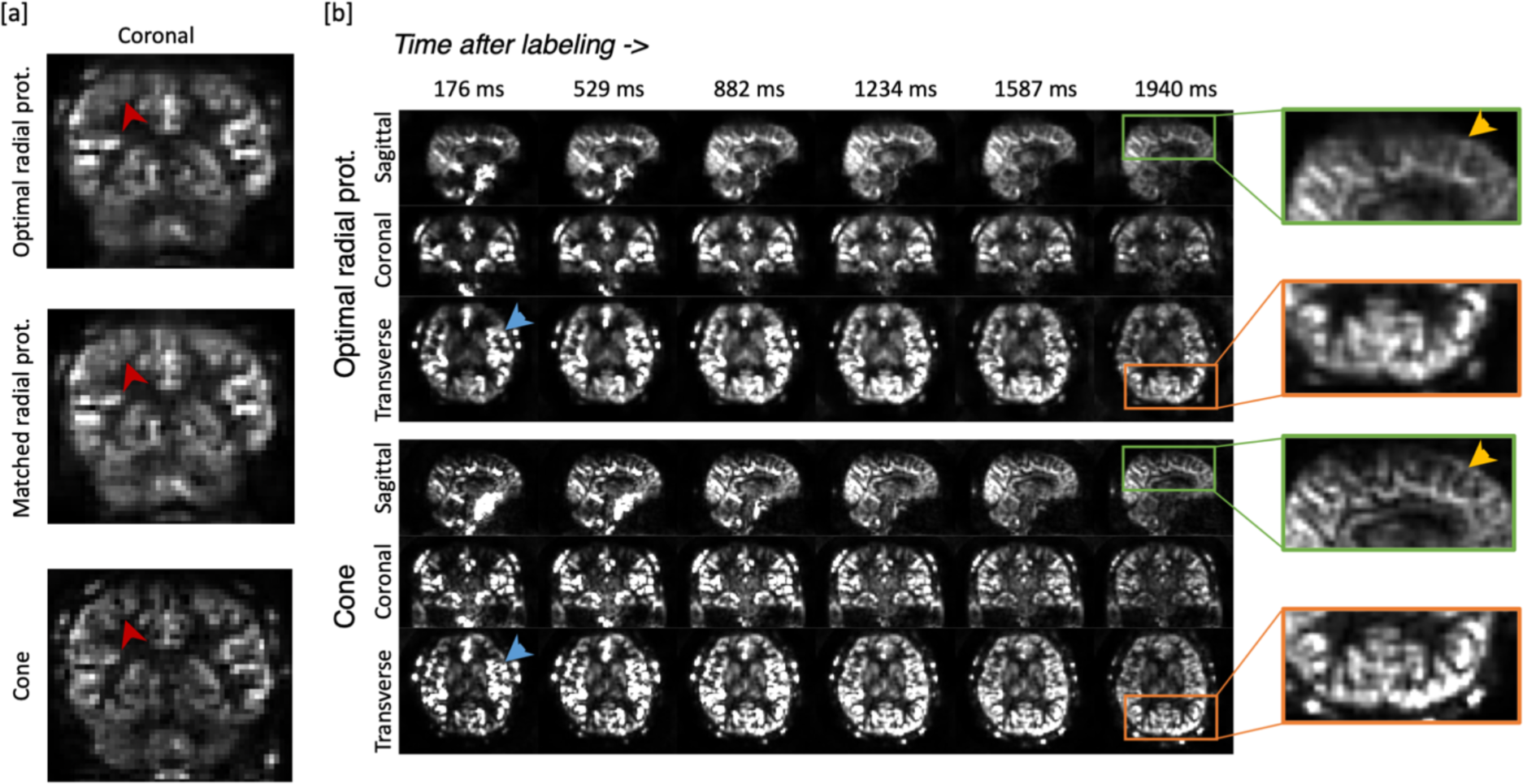
Example perfusion images acquired using different trajectories. [a] Coronal slice of an example frame for each of the three protocols, demonstrating the improved spatial resolution achieved with the proposed cone trajectory (red arrowhead). [b] Comparison between the proposed cone trajectory and the better-performing radial trajectory (using the “optimal radial protocol”^6^) showing example sagittal, coronal and transverse slices for all six frames. The displaying window of each frame is rescaled to its maximum intensity for better clarity. Zoomed sections again demonstrate the improved spatial resolution achievable with the cones approach.

Figure 5b compares radial and cone data across all the time points in multiple orientations. The initial timepoints showed the clustering of high-intensity signals, which could be attributed to the presence of labeled blood within vessels. Specifically, vessels marked by the blue arrow exhibited clear contours in the cone trajectory results, whereas the radial counterparts displayed noticeable blurring. The final frame corresponds to a typical Post-Labeling Delay (PLD) used in single timepoint protocols^35^, where large vessel signal is generally absent in healthy volunteers. Enlarged regions in the green and orange boxes highlight resolution improvements also in this perfusion-weighted signal using the cone trajectory.

### In-vivo angiography results

Figure 6 presents a visual comparison of in-vivo angiography results obtained from the same raw datasets as the aforementioned perfusion images, albeit reconstructed at a higher spatial and temporal resolution.

**Figure 6.**
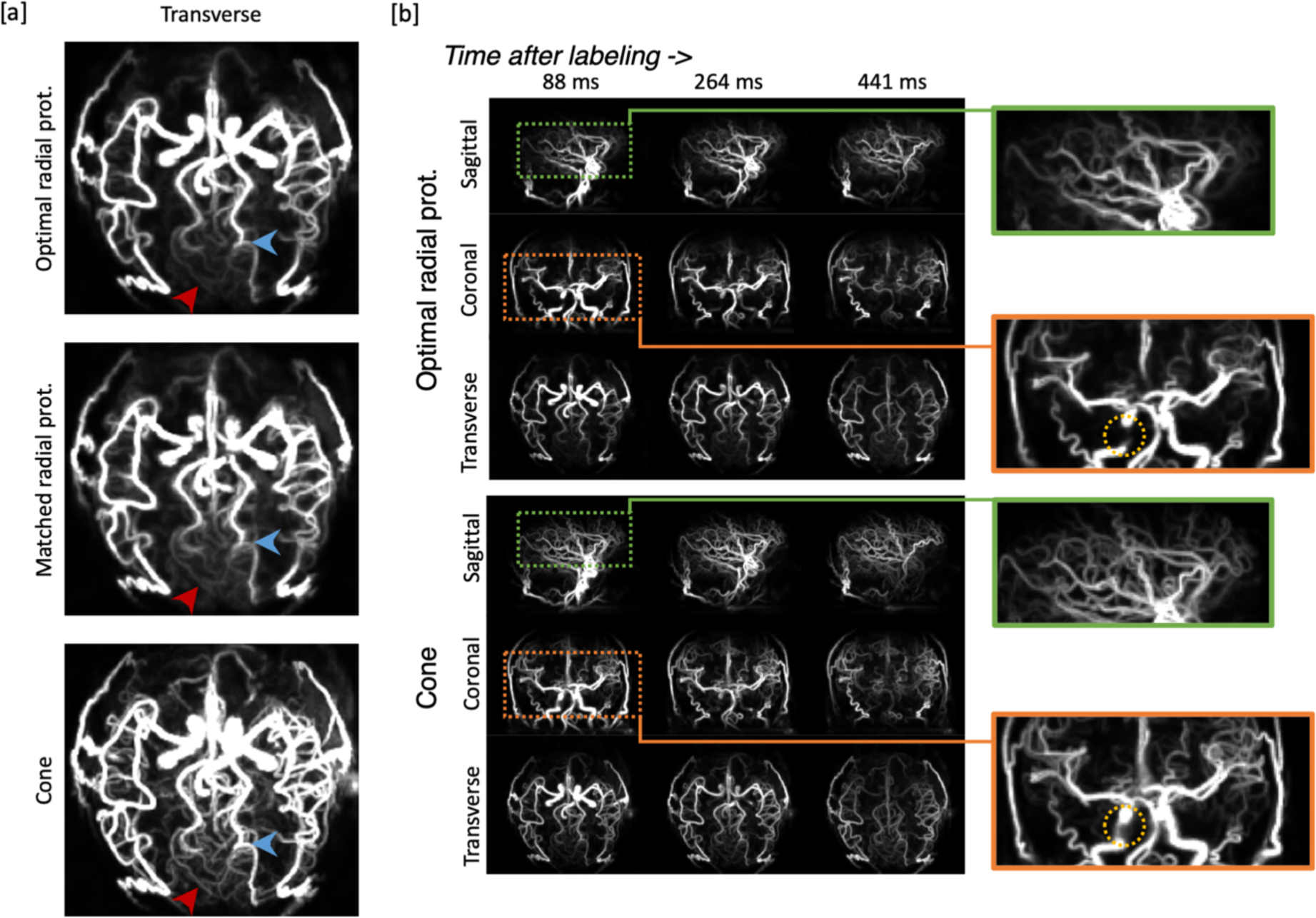
Example angiographic images acquired using different trajectories. [a] transverse maximum intensity projection of an example frame in angiographic images reconstructed using the same set of raw data as the perfusion images in Figure 5. The improved visualization of small vessels (red arrows) emphasizes the superior resolution of the proposed cone trajectory. [b] Comparison of angiographic images between radial and cone trajectories (Maximum Intensity Projections) in three example frames. Two regions were enlarged for both trajectories respectively. Orange box: The cone trajectory provided better visualization of the proximal arteries, reducing signal dropout due to B_0_ inhomogeneity and/or flow effects. Green box: The cone trajectory allowed clearer visualization of small distal vessels than the radial trajectory.

In Figure 6a, three illustrative transverse view maximum intensity projections acquired with the three different protocols are presented. Clear vessel delineation was achieved in all cases with clean subtraction of background tissue. However, focusing on the vessels highlighted by the red arrow, the cone protocol provided a much better visualization of the small distal vessels due to its improved spatial resolution resulting from the better coverage of peripheral k-space. Of the two radial protocols, it is evident that the radial trajectory with TR matched to the cone protocol yields marginally blurrier outcomes compared to the radial trajectory with the previously optimized protocol, which is expected due to the smaller number of spokes acquired. Notably, the vessel’s sharpness, indicated by the blue arrow, is diminished in the former.

To facilitate clarity, the comparison between the cone trajectory and the optimal radial protocol is presented in Figure 6b, showcasing three exemplary frames for each protocol in all three views. The advantage of the cone trajectory lies in the enhanced visualization of distal vessels with sharper contours, consequently improving the diagnostic and vessel-related analysis capabilities. In the orange box, particularly within the region enclosed by the yellow circle, a conspicuous disparity emerges: the radial approach fails to preserve signal intensity within some segments of the proximal arteries, likely attributed to signal dephasing due to B_0_ inhomogeneity and/or flow effects. In contrast, the cone trajectory has less signal loss, thanks to its center-out design and substantially shorter echo time.

### Quantitative comparisons

The quantitative repeatability evaluation of the perfusion images is presented in Figure 7a. The observed trend in repeatability follows a gradual decline as the post-labeling delay increases, reflecting the signal’s natural attenuation after labeling. In agreement with the visual comparisons in Figure 5, the cone trajectory was more repeatable than the two radial trajectories. Similar result for the angiographic images is shown in Figure 7b. Given the considerably stronger angiography signal compared to perfusion, the absolute values of scan repeatability were inherently higher. In both radial trajectories’ results, repeatability declined quickly as frames progressed. In contrast, the cone trajectory maintained a relatively higher repeatability across frames. The consistency further substantiates the robustness and reliability of our proposed cone design. The significance for the cone trajectory improvement was confirmed by multi-way ANOVA test (p<0.001).

**Figure 7.**
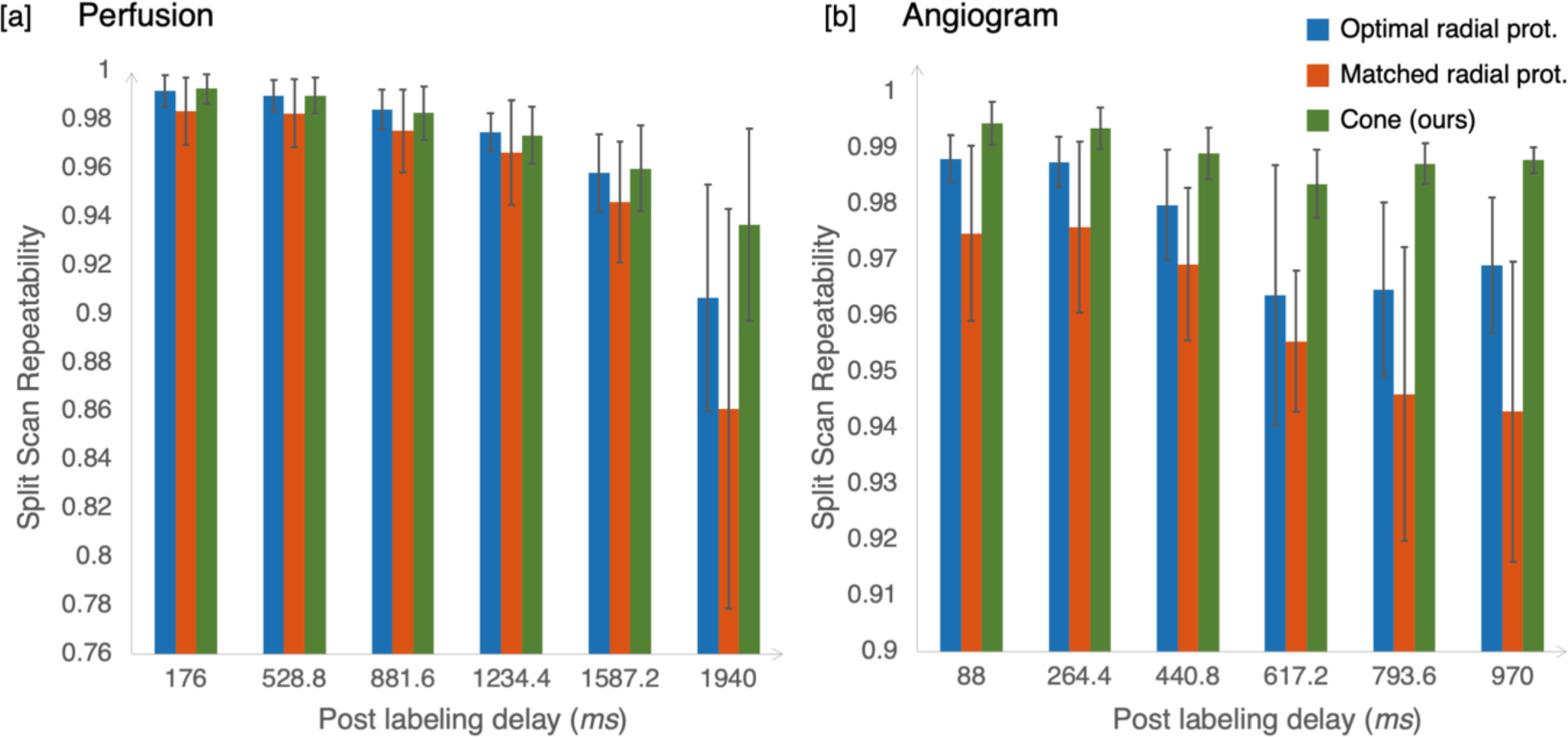
A) Comparison of the repeatability of the perfusion images between three protocols: 1) the optimal radial protocol, 2) the matched radial protocol, and 3) our cone trajectory. The mean and standard deviation across subjects are shown here. Overall, the scan repeatability decreased with the PLD due to the signal attenuation. The cone trajectory had the highest overall scan-rescan repeatability of three protocols, particularly at later PLDs, and its performance was the most stable across different subjects (*p* < 0.001 in multi-way ANOVA test). B) Comparison of the repeatability of the angiographic images between the three protocols. Only the first 6 frames out of the 12 reconstructed frames were used for repeatability calculation, as later frames are dominated by perfusion signal and the vessel mask was no longer applicable. Since the angiography signal is much stronger than perfusion, the average scan repeatability is higher than perfusion. However, the cone trajectory still maintains the best and most stable performance across different subjects and frames, and the significance was confirmed by multi-way ANOVA test (*p* < 0.001) as well> 0.001 in multi-way ANOVA test). B) Comparison of the repeatability of the angiographic images between the three protocols. Only the first 6 frames out of the 12 reconstructed frames were used for repeatability calculation, as later frames are dominated by perfusion signal and the vessel mask was no longer applicable. Since the angiography signal is much stronger than perfusion, the average scan repeatability is higher than perfusion. However, the cone trajectory still maintains the best and most stable performance across different subjects and frames, and the significance was confirmed by multi-way ANOVA test (*p* < 0.001) as well.

## Discussion

In this work, a novel two-stage curved cone trajectory design was proposed and integrated into the 4D CAPRIA sequence to improve the spatial resolution and SNR of both the angiographic and perfusion images. This allowed high quality dynamic 3D maps of blood flow through the vessels and into the tissue bed to be acquired simultaneously in a single short scan (6 mins), providing a comprehensive evaluation of cerebral blood flow. A process using numerical simulations and physical phantom acquisitions for optimizing the cone design parameters suitable for our application was presented. LLR regularization was used within the in vivo reconstruction to take advantage of the incoherent sampling of our trajectory and reduce artifacts and noise. The cone trajectory was compared against the previously optimized 4D CAPRIA radial trajectory on a group of healthy subjects, demonstrating that our cone design could improve 4D CAPRIA through higher spatial resolution, improved scan-rescan repeatability, and reduced signal loss in proximal arteries.

The most straightforward explanation for the improvement from radial to our cone design is the increased sampling efficiency and better utilization of the gradient system. The radial trajectory features oversampling at the k-space center, but it does not make full use of the degrees of freedom afforded by the gradients. Our proposed cone trajectory design is based on the intuition that traversing more of k-space in each readout by adding twists to the radial spokes will be beneficial both to improving the resulting point spread function by improving sampling in peripheral k-space, as well as more uniform sampling throughout k-space which is beneficial for SNR ^36^. A large cone angle 60° was also adopted, as we speculate that trajectories with larger cone angles has a larger gradient perpendicular to radius and traverse longer distance at the k-space center compared to those with smaller cone angles and thus more robust to signal variations. It agreed with the observations from the phantom scanning, which, in a given readout time, cone trajectories with larger angles exhibit less aliasing due to signal variations.

Additional flexibility in the design of the trajectory over existing approaches was desired for better adaptation to our combined imaging of two modalities with different spatiotemporal resolution requirements. To be more specific, if only angiography is considered, the time spent at the k-space central region is less important than focusing on traversing quickly to peripheral k-space. Therefore, a two-stage design was introduced, which allowed for different trajectory properties in central and peripheral k-space to better accommodate both modalities. Additional simulated results comparing the proposed two-stage design with existing base cone trajectories are shown in Supplementary Figure 7, demonstrating the smaller FWHM for angiography and higher effective SNR for both angiography and perfusion reconstructions using our cone design, compared to Johnson’s^13^ and Gurney’s^9^, when combined with the golden ratio rotation approach. Because of the complex interaction between signal variations, off-resonance sensitivity, and non-Cartesian undersampling artifacts, a purely simulation-based optimization would have been unreliable.

Instead, a two-step process combining numerical simulation and physical phantom acquisition was adopted to find suitable parameters for the proposed cone trajectory. To ensure a trajectory versatile to variable spatial and temporal resolution, advanced non-linear reconstruction method was not used in searching for optimal parameters. However, the improvement of PSF and reduction in artifacts observed with vanilla reconstruction would also transfer to more advanced reconstruction method. The optimization not only covered a considerable range of possible values but also reflected realistic scanning conditions faithfully. One exception was the parameter *m*, which was heuristically chosen by placing the separation point at the middle of two parts of the cone trajectory. Preliminary experiments varying *m* found the it interacted with other parameters during the optimization process, therefore *m* was fixed for two-stage cone trajectory to reduce complexity.

Our design also possesses a range of advantages when compared with other non-Cartesian trajectories developed over the years. Conventionally, non-Cartesian trajectory design focused on the fully sampled case, where rigorous calculation is performed to satisfy the Nyquist sampling theorem, from 2D spiral^8,37^ to 3D cone.^9,11^ Given careful analytical formulations, these trajectories are fast to compute but cannot be easily adapted to a more flexible highly undersampled case, which we required to achieve clinically feasible acquisition time for this application. The approach taken in the original radial-cones^13^ trajectory did not require precise adherence to the Nyquist sampling requirements, but the design and optimization were still targeted to achieve approximately full sampling. Our design, however, has a set of free parameters which can be determined through a two-step process to be fine-tuned for specific applications without the requirement for full sampling. Additionally, as our trajectory is generated within the sequence code, the trajectory parameters can be easily adjusted during scanning without manually loading a new trajectory, as with numerically optimized trajectories^15,16^. In addition, the use of golden ratio rotations means the undersampling factor and temporal resolution can be chosen arbitrarily during reconstruction. For example, supplementary Figure 11 demonstrates exemplary reconstructions using various temporal resolutions.

Despite the improved sampling efficiency, the additional twists in the cone trajectory design make the acquisition more susceptible to gradient imperfections such as gradient delays. The mismatch between the actual and expected trajectory can introduce errors into the reconstruction. For our trajectory, the slew rate and gradient were limited to reduce the effect of the trajectory mismatch. No notable gradient imperfection related artifacts were observed in all our experiments. However, use of the gradient impulse function^38,39^ could improve the reconstruction further and allow the full potential of the gradient system to be utilized.

In-vivo scans in clinical settings often suffer from patient motion. Cone design maintains the ability to use retrospective motion correction while the improved sampling at the central k-space also enables better quality navigator for more accurate registration. In this preliminary investigation, a small group of 8 relatively young healthy subjects were scanned. In the future, a broader comparison within a larger group of older healthy subjects and patients with cerebrovascular disease should be carried out for a more comprehensive validation of the generalizability of our cone design. We could also perform a thorough comparison with more conventional methods and explore the quantification of blood flow from these images^40,41^.

### Conclusion

In this work, we introduced an innovative 3D cone trajectory design, combined with a golden ratio rotation strategy, to enhance SNR and resolution without prolonging acquisition time. Our design better utilizes the gradient system, covering more k-space than the radial trajectory in a single readout. Notably, its non-Cartesian artifacts exhibit enhanced incoherence and uniformity compared to the radial approach, making it more suitable for LLR reconstruction. Through a comprehensive process involving numerical simulations and physical phantom scans, a combination of the most suitable trajectory parameters was selected. The evaluation on a group of healthy subjects demonstrated that our cone trajectory surpassed the original 4D CAPRIA approach, not only in terms of spatial resolution but also in scanning repeatability for both angiography and perfusion images.

## DATA AVAILABILITY STATEMENT

The code for generating the trajectory and aforementioned simulation is openly available on Github (https://github.com/Michaelsqj/TrajectoryDesign). The code used for image reconstruction used in the study can be found at (https://github.com/Michaelsqj/cone_recon). Deidentified image data will be made available on the WIN Open Data server in the future. This is currently in development.

## Acknowledgements

We are grateful for funding support from a Sir Henry Dale Fellowship jointly funded by the Wellcome Trust and the Royal Society (220204/Z/20/Z). The Wellcome Centre for Integrative Neuroimaging is supported by core funding from the Wellcome Trust (203139/Z/16/Z) with additional support from the NIHR Oxford Health Biomedical Research Centre (NIHR203316). W.W. is supported by the Royal Academy of Engineering (RF\201819\18\92). MC is supported by the Canada Research Chairs program. The views expressed are those of the authors and not necessarily those of the NIHR or the Department of Health and Social Care. Many thanks also to Jeff Fessler, Philipp Ehses and colleagues for making available their excellent NUFFT and Siemens raw data reading MATLAB code, as well as to Siemens Healthineers for providing the base pulse sequence code, that we built upon in this work. For the purpose of open access, the author has applied a CC BY public copyright license to any Author Accepted Manuscript version arising from this submission.

