## Supplementary Figure for "Efficient 3D cone trajectory design for improved combined angiographic and perfusion imaging using arterial spin labeling"

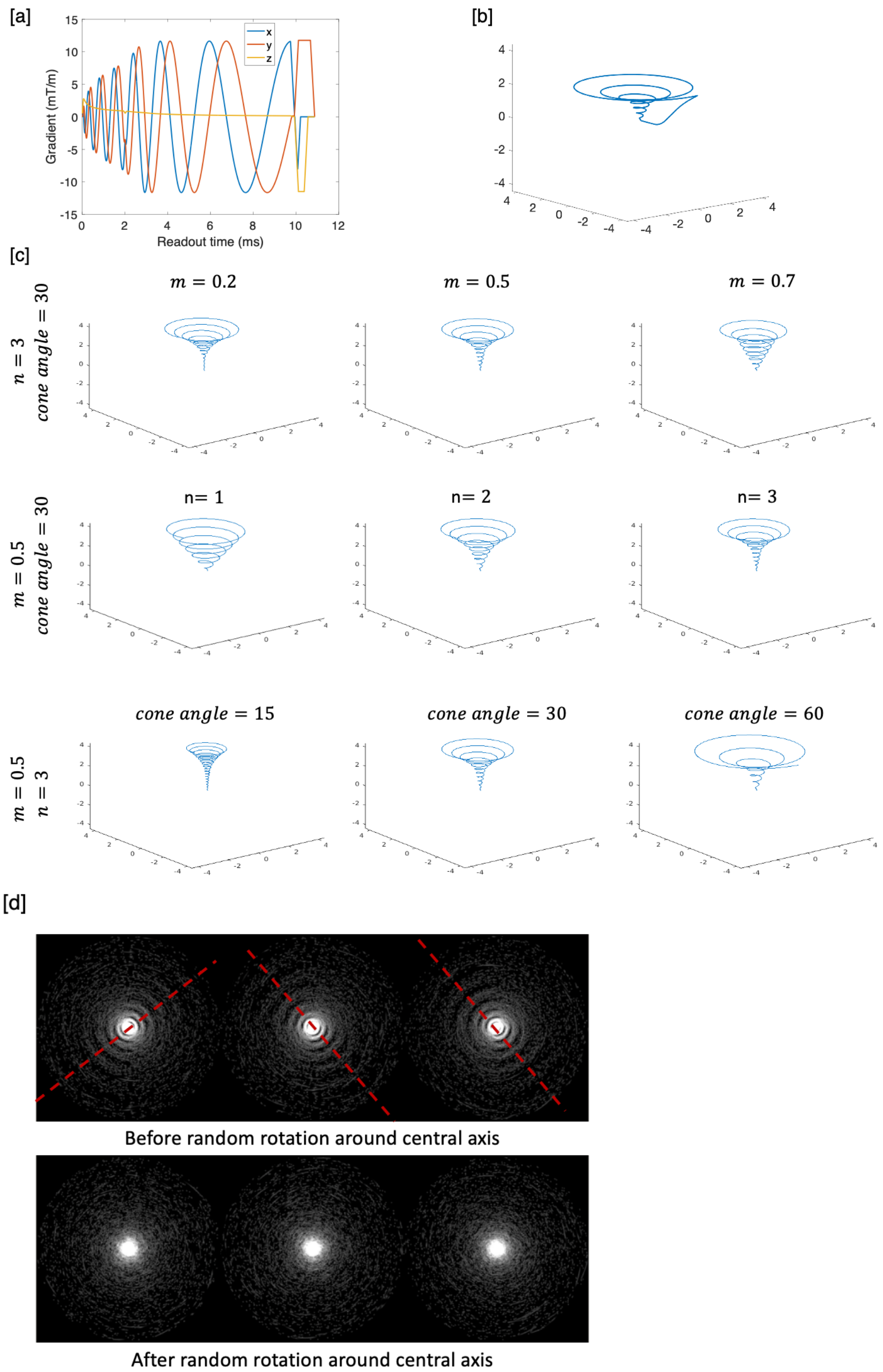


Figure S1. A) Example gradient waveforms. B) corresponding trajectory in k-space. C) The effect of varying the free parameters in the proposed cone design on the trajectory shape. Fixed parameters are displayed on the left of each row, then variation in one other parameter is shown: m (top row), n (middle row) and the cone angle, θ (bottom row). D) More uniform sampling in k-space using random rotation around the central axis of each shot.

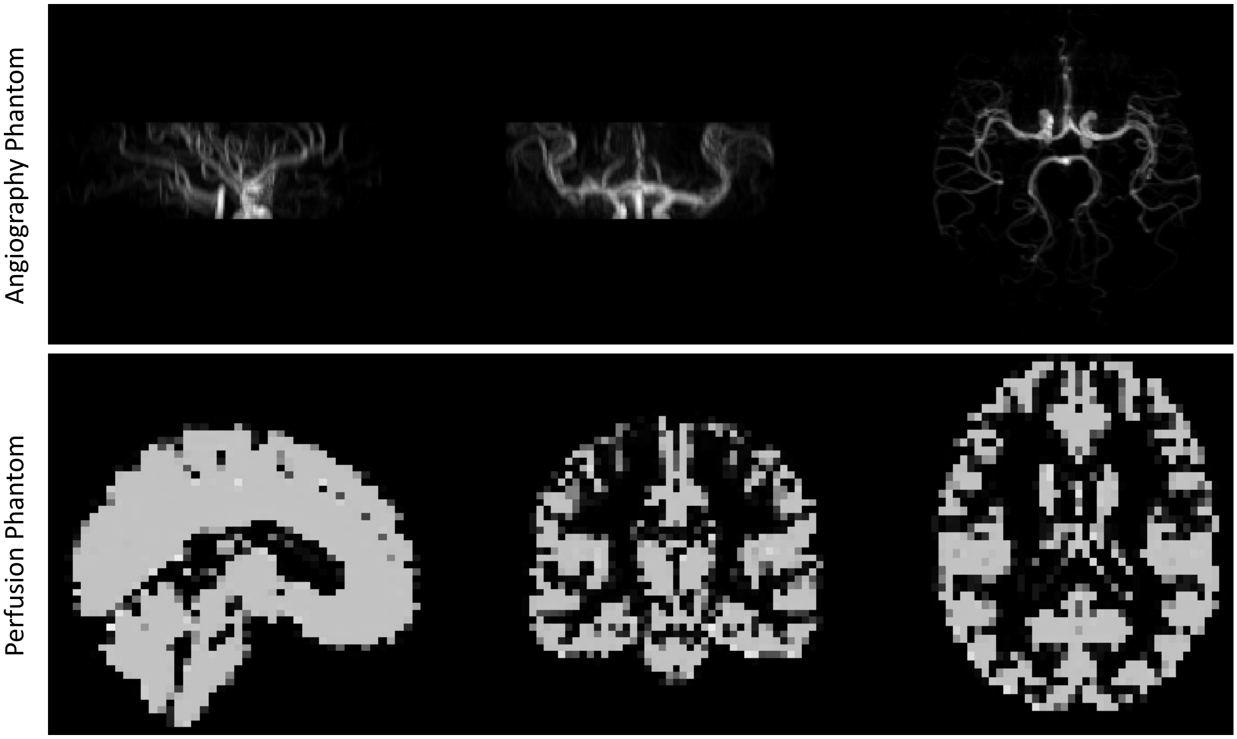


Figure S2. Numerical phantoms for angiography (shown as maximum intensity projections, top) and perfusion imaging (example slices in all three planes, bottom) that were used for simulation experiments.

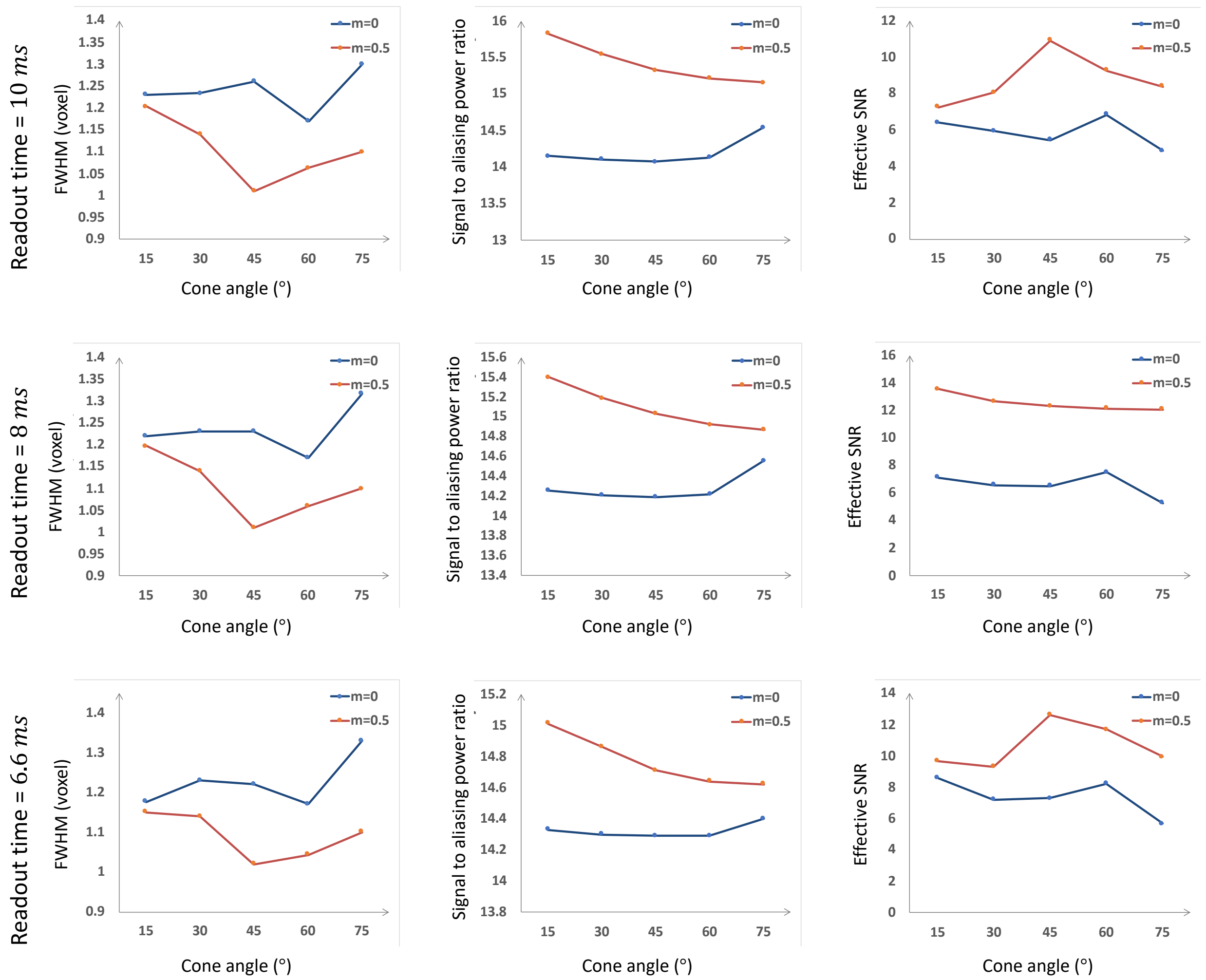


Figure S3. Numerical simulations for angiographic imaging, showing variation in PSF FWHM (left), signal to aliasing power ratio (middle) and effective SNR (right) with variable readout times (top to bottom), m values and cone angles. In these simulations, n was fixed at 3.

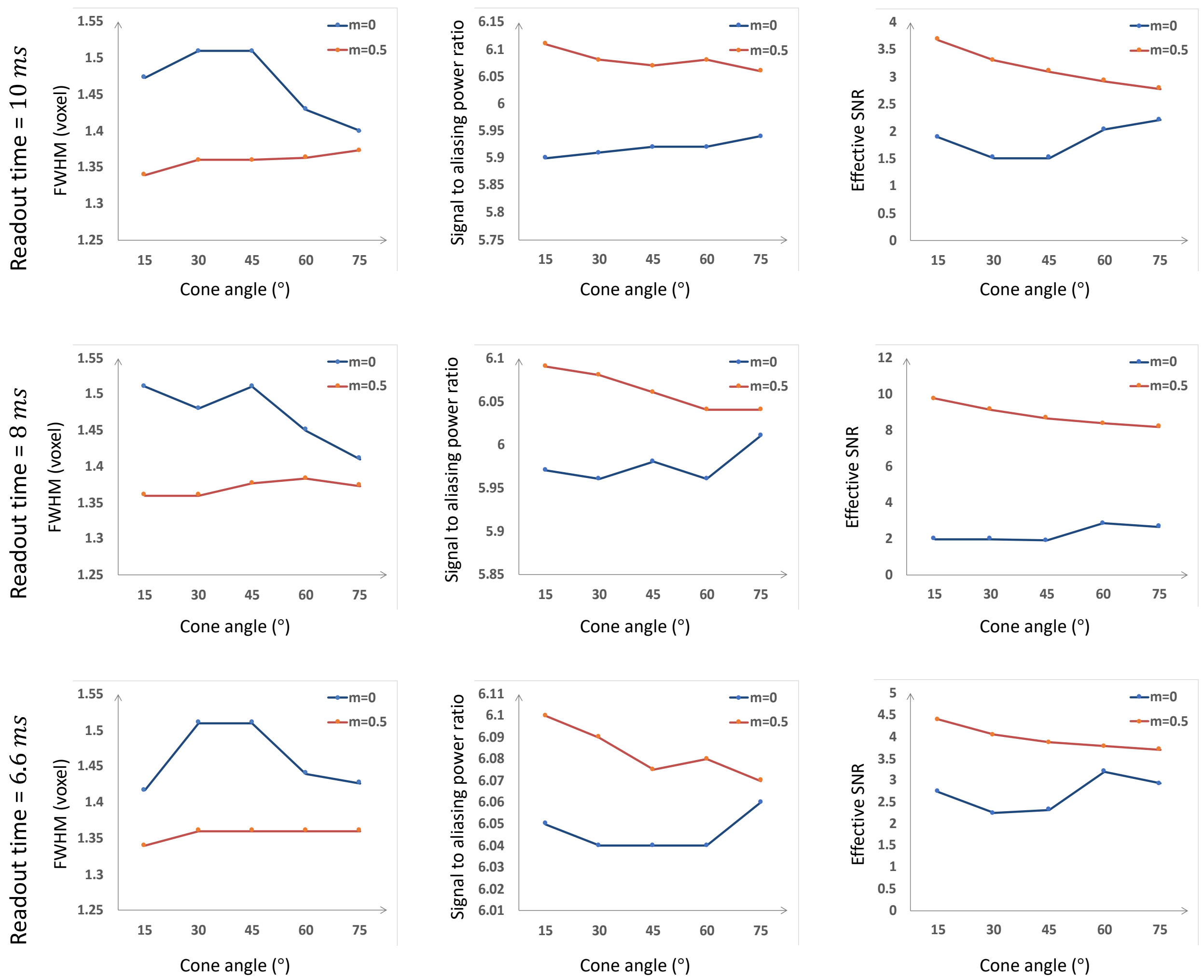


Figure S4. Numerical simulations for perfusion imaging, laid out as in Figure S3.

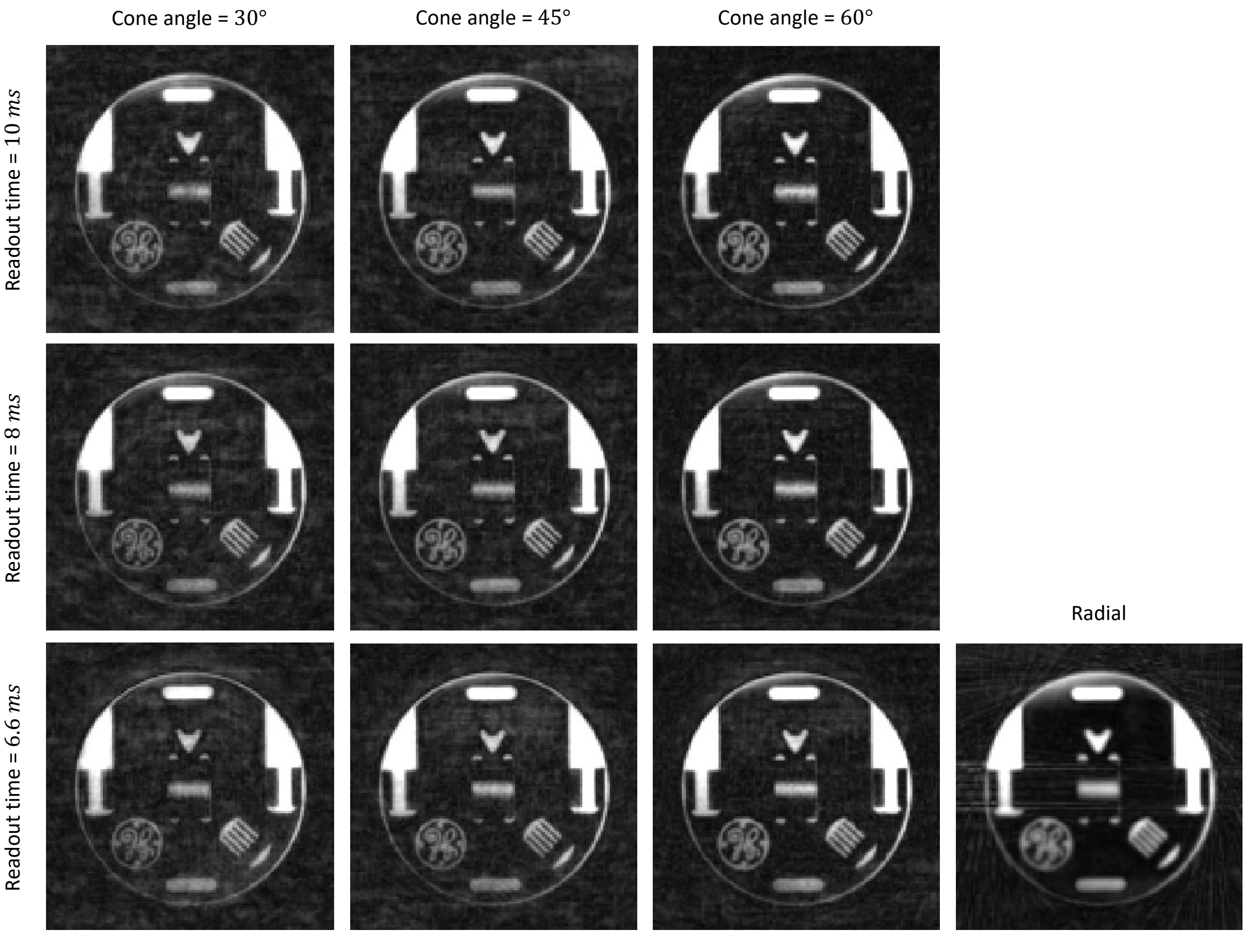


Figure S5. Physical phantom comparison, view #1.

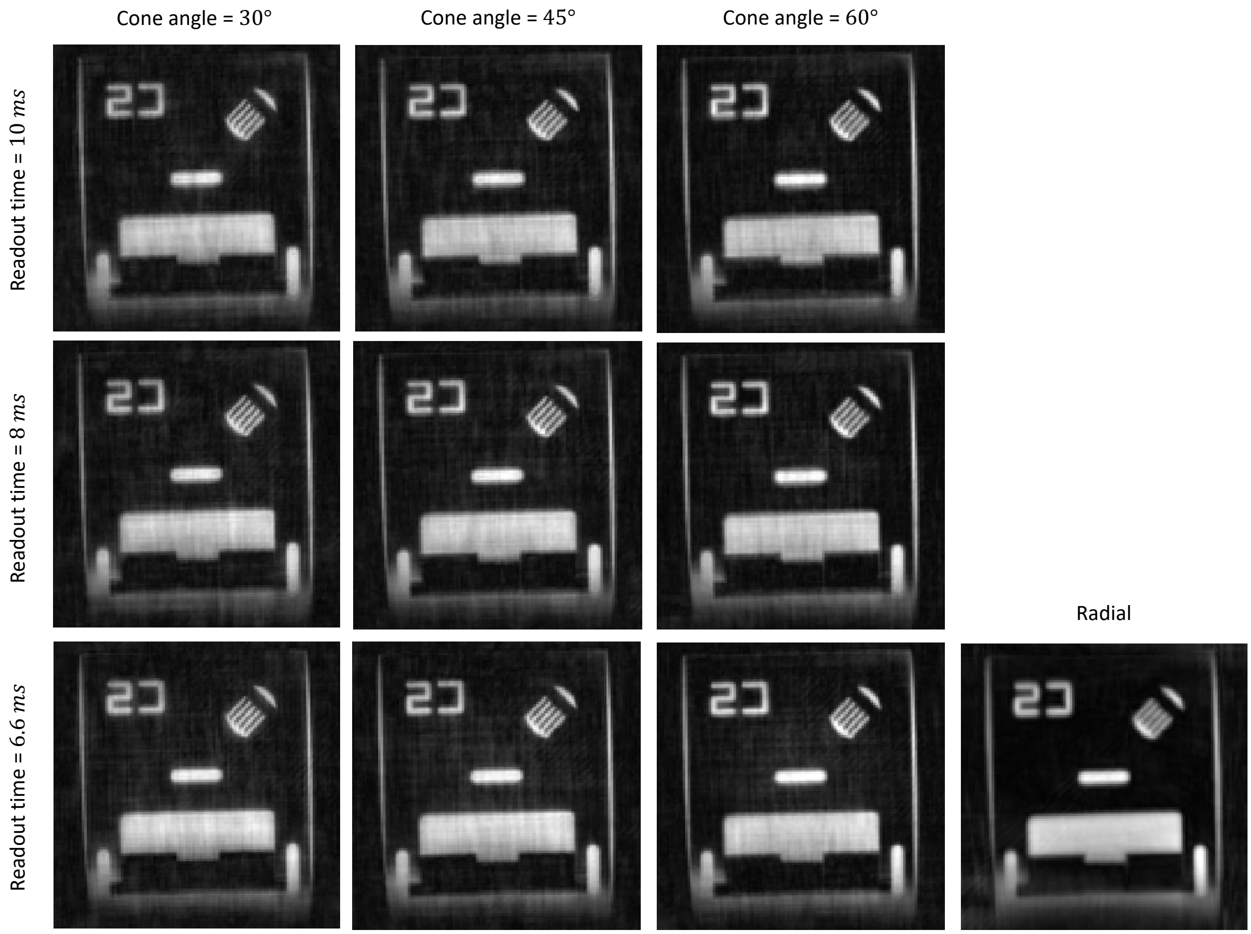


Figure S6. Physical phantom comparison, view #2.

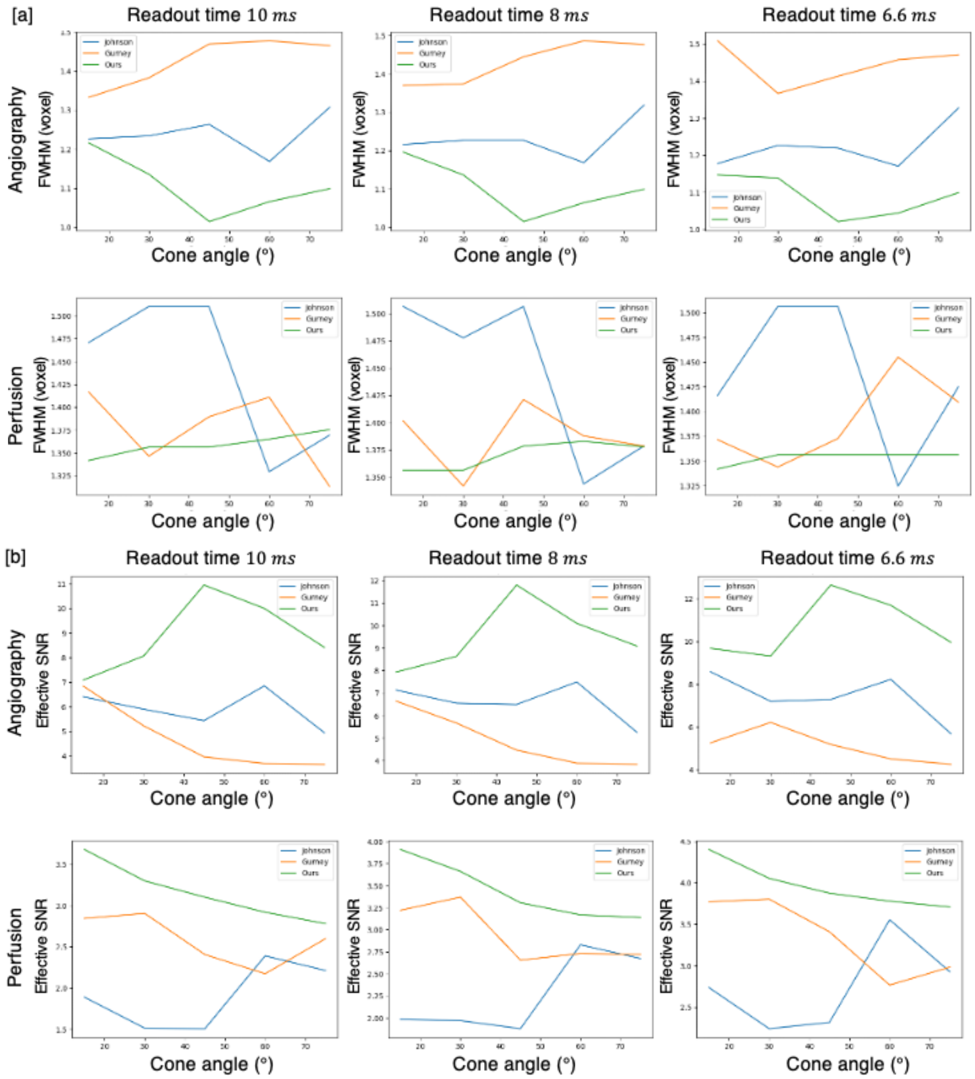


Figure S7. Simulated comparison with Gurney's[1] cone and Johnson's[2] cone in both angiogram and perfusion

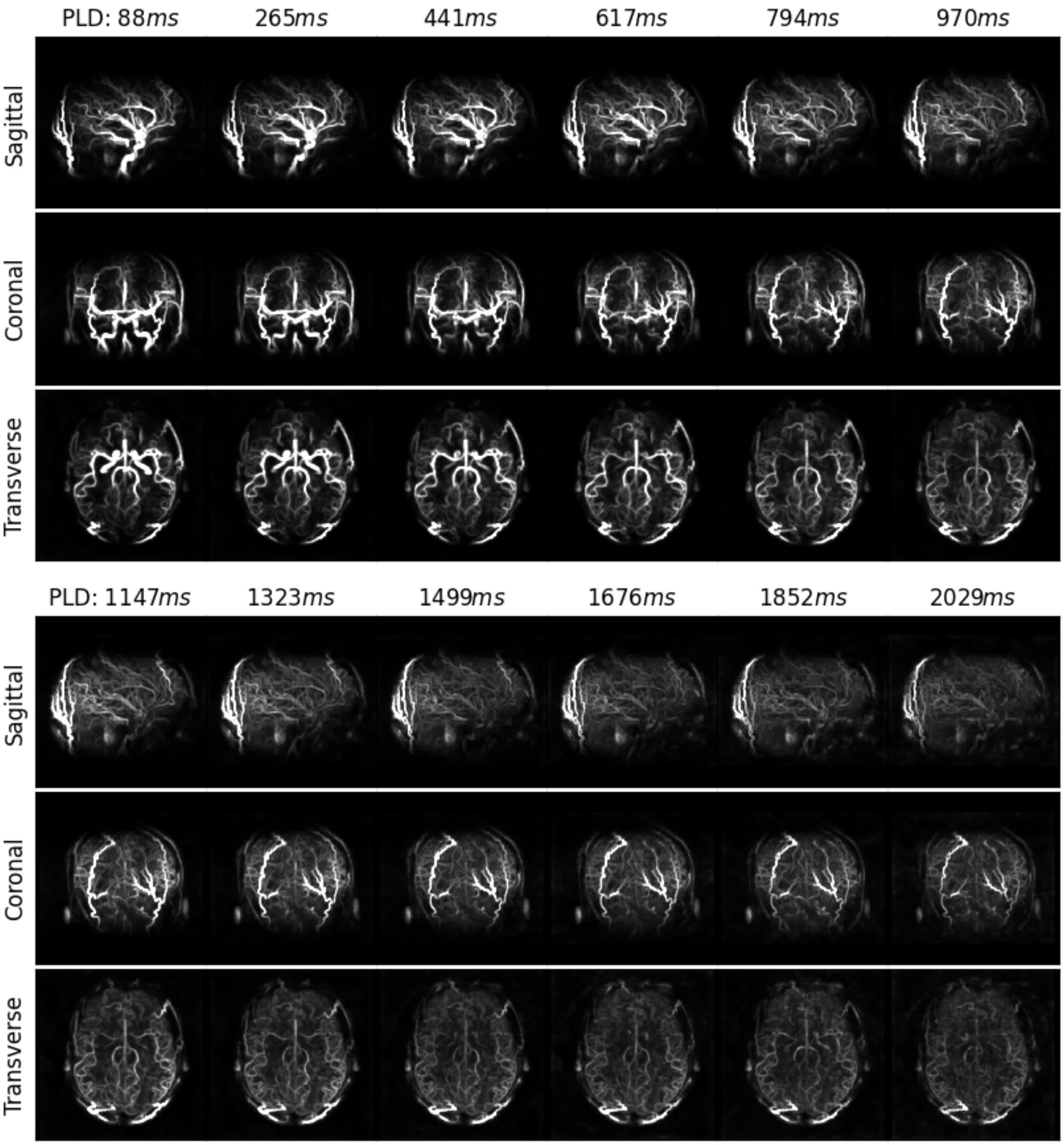


Figure S8. All reconstructed angiography frames in one subject acquired with the 4D CAPRIA protocol using a radial trajectory.

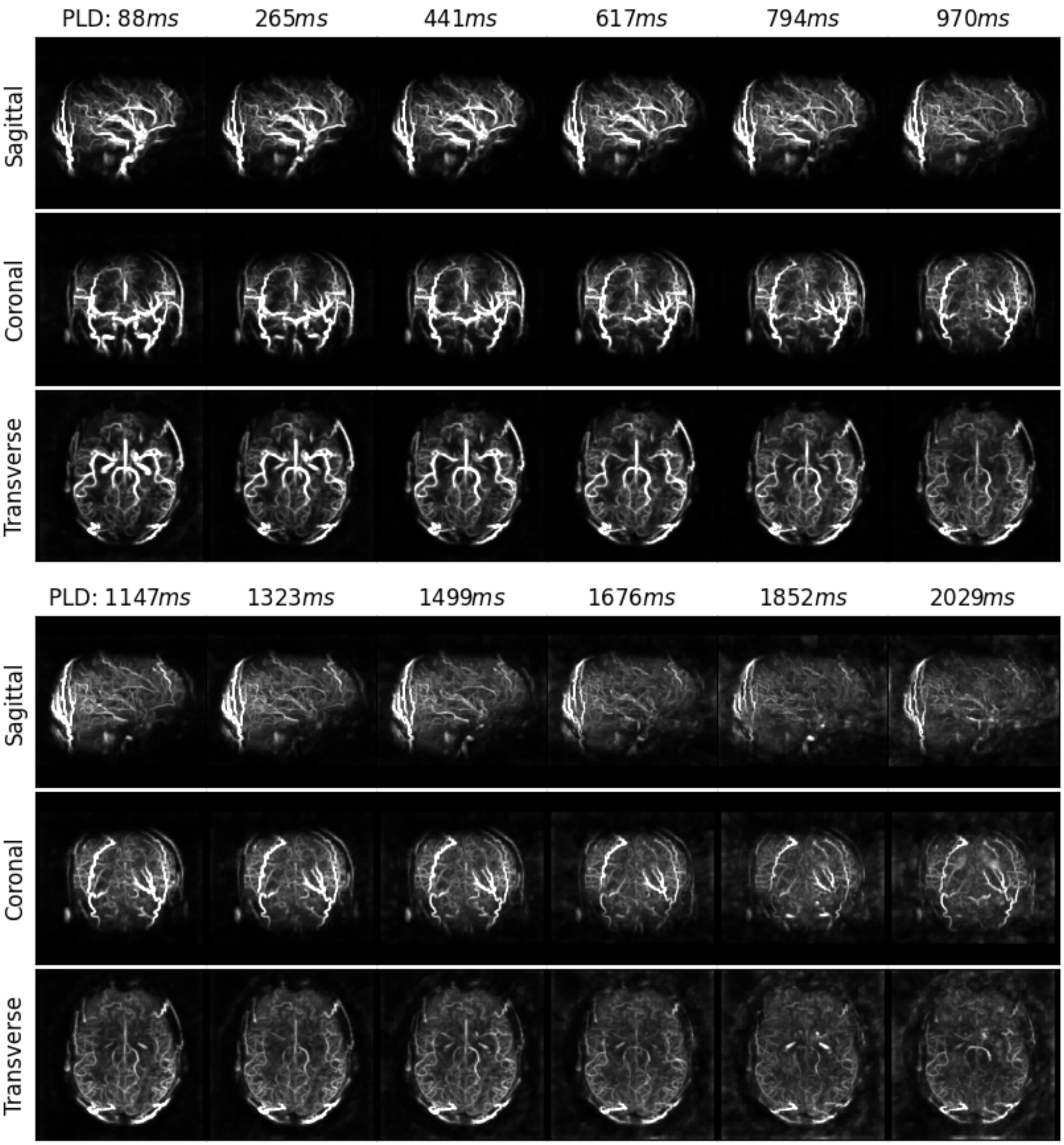


Figure S9. All frames of angiography image acquired by radial trajectory (TR matched with cone protocol)

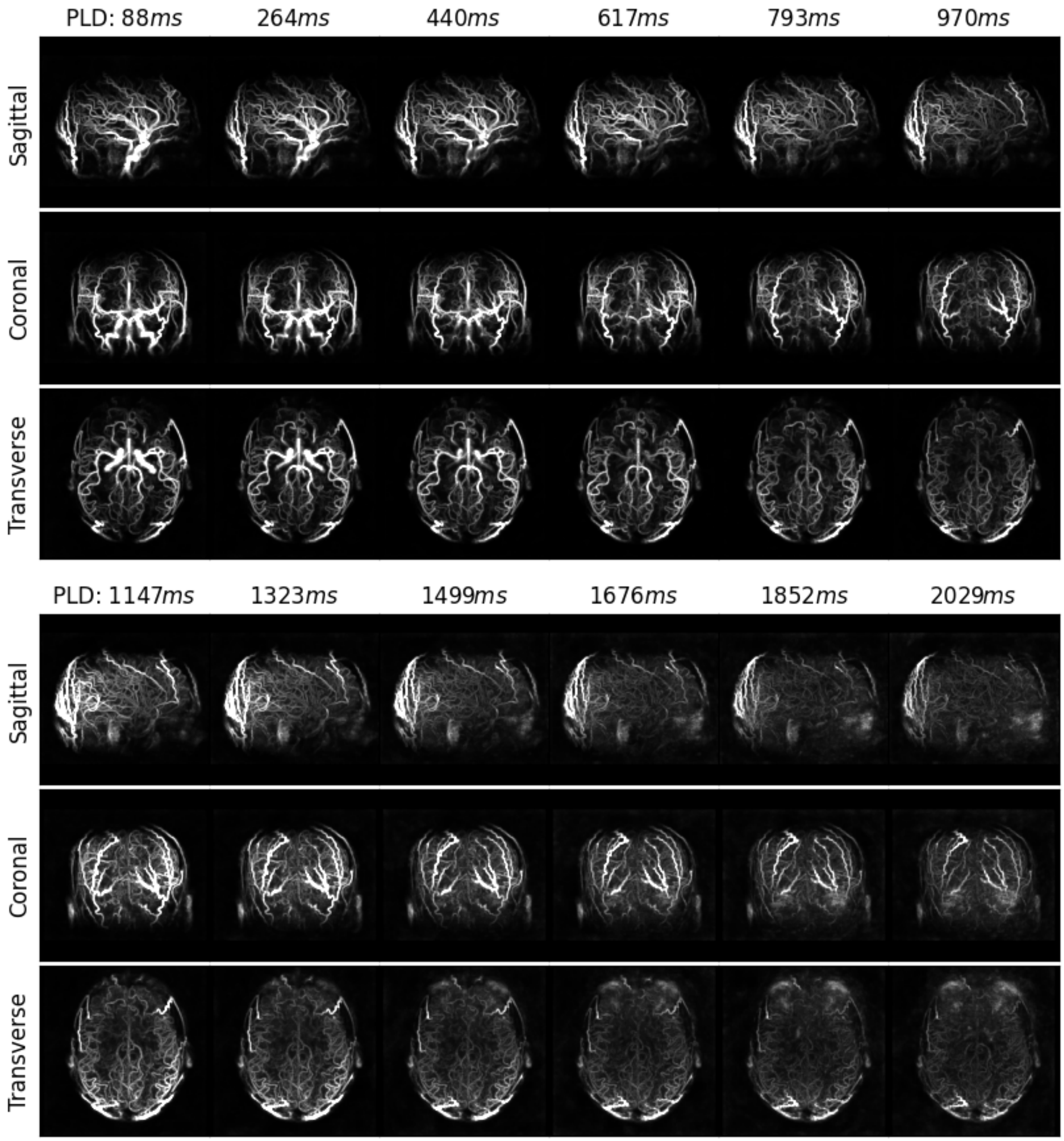


Figure S10. All frames of angiography image acquired by the cone trajectory

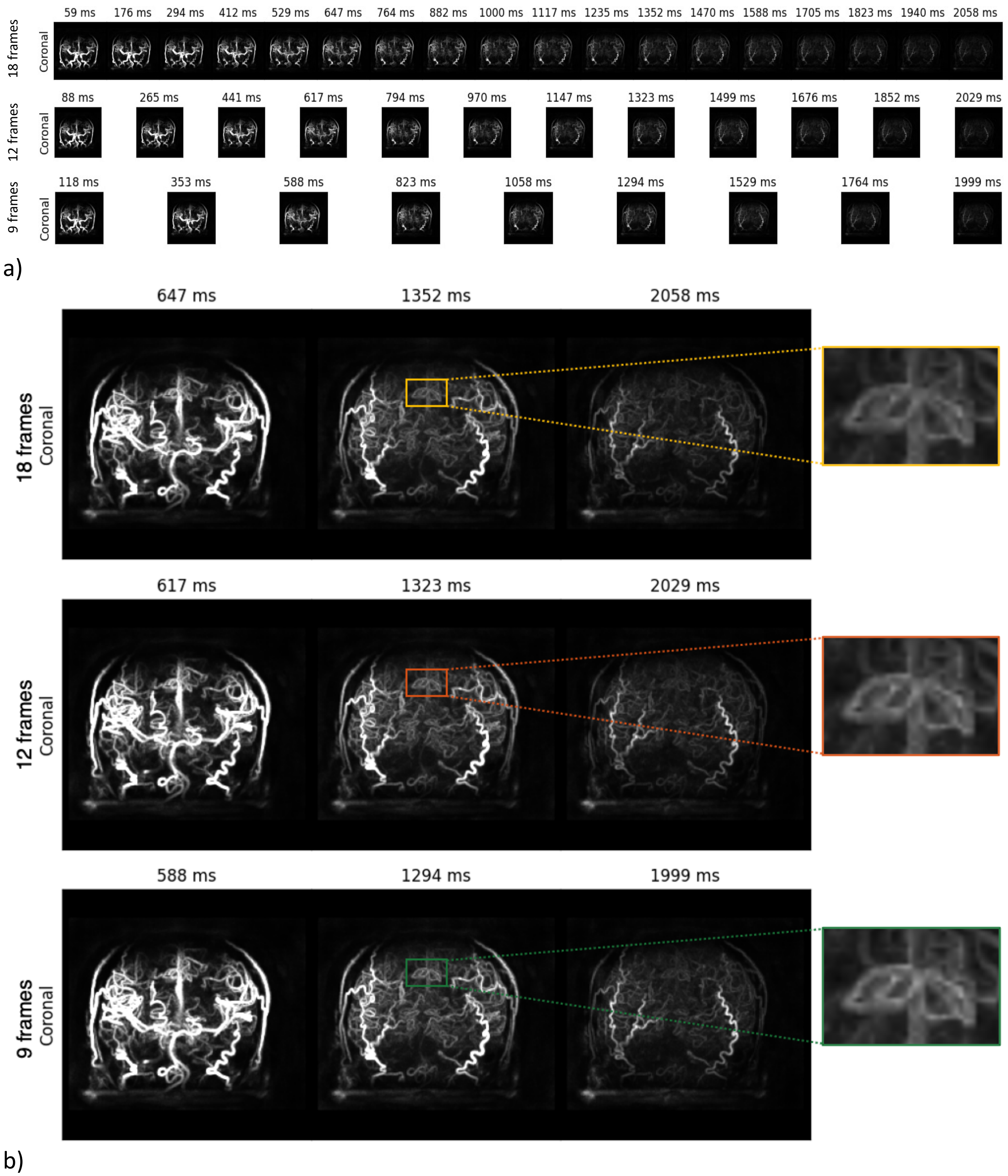


Figure S11. A) CAPRIA sequence using cone trajectory and 3D Golden Angle enables flexible reconstruction of dynamic angiogram with different temporal resolution. B) There is a slight tradeoff between temporal resolution and spatial resolution. As shown in the zoomed-in boxes, vessels in the 9-frame result are sharper than those in the 18-frame one.

1. Gurney PT, Hargreaves BA, Nishimura DG. Design and analysis of a practical 3D cones trajectory. *Magn Reson Med*. 2006;55(3):575-582. doi:10.1002/mrm.20796

2. Johnson KM. Hybrid radial-cones trajectory for accelerated MRI: Hybrid Radial-Cones Trajectory for Accelerated MRI. *Magn Reson Med*. 2017;77(3):1068-1081. doi:10.1002/mrm.26188
